# Temporal Structure of Reward Availability and Sensory Uncertainty Modulate Allocation Dynamics in Naturalistic Foraging

**DOI:** 10.64898/2026.04.14.718537

**Authors:** Panos Alefantis, Yaxin Guo, Noushin Quazi, Cristina Savin, Dora E. Angelaki, Xaq Pitkow, Najib J. Majaj

**Affiliations:** Department of Psychology, New York University, New York, NY 10003, USA; Center for Neural Science, New York University, New York, NY 10003, USA; Department of Biomedical Engineering, NYU Tandon School of Engineering, New York, NY 11201, USA; Carnegie Mellon University, Pittsburgh, PA, USA; Center for Data Science, New York University, New York, NY 10011, USA; Tandon School of Engineering, New York University, New York, NY 11201, USA

## Abstract

Adaptive foraging requires animals to combine uncertain sensory cues with predictions about when rewards are likely to occur. While theoretical models describe how animals should allocate their effort under variable-interval reward schedules, it remains unclear how the timing or rewards and the reliability of sensory cues affects behavior. We developed a continuous foraging task in which freely moving macaque monkeys navigated among three reward patches. Rewards became available at unpredictable times, with their availability signaled by a visual cue of varying reliability. We also varied the schedule of reward availability: in some conditions, rewards were equally likely to become available at any moment (exponentially distributed intervals), while in others the interval distribution was more concentrated around a particular mean (gamma-distributed intervals) which increased the cost of premature responses. Under exponential schedules, monkeys eventually allocated their time at different patches according to reward schedules, and cue reliability had only modest effects. Under gamma-distributed intervals, monkeys more quickly learned to differentiate between patches. Their choices were more strongly dependent on predicted reward timing, particularly when sensory cues were highly reliable. These results show that both the timing of rewards and reliability of sensory cues shape how animals allocate their time and effort in continuous naturalistic foraging tasks.

## Introduction

Foraging is among the most fundamental behaviors across living organisms. When choosing among multiple food sources, animals distribute their choices across options to maximize total reward intake, typically guided by reward probability and timing. As environmental conditions change, effective foraging requires animals to continuously update their estimates of reward availability from experience and current sensory evidence.

Decades of work on concurrent variable-interval schedules have established core behavioral phenomena of reward-based allocation. When animals choose between simultaneously available options that deliver rewards at different rates, they distribute their choices approximately in proportion to relative reward rates, a regularity formalized as the matching law (Herrnstein, 1961; Houston et al., 2021). This law has been demonstrated across species, reinforcer types, and procedural variations (Baum, 1974; Davison & McCarthy, 1988), establishing it as one of the most robust regularities in the study of choice behavior.

Neurophysiological studies in non-human primates have identified neural substrates underlying how subjects distribute their choices in proportion to reward rates. Monkeys trained on concurrent variable-interval schedules exhibit reward-proportional allocation, and neurons in lateral intraparietal cortex (LIP) and anterior cingulate cortex (ACC) encode relative value, experienced reward rate, and opportunity cost (Sugrue et al., 2004; Corrado et al., 2005; Kolling et al., 2012; Hayden et al., 2011; Blanchard et al., 2015). Yet the paradigms used in these studies impose constraints that deviate from the foraging problems animals actually face. Subjects are head-fixed, choices are expressed through eye movements between two spatially fixed targets, and reward schedules are presented in discrete trials in which stimulus conditions are fixed within each trial. Crucially, in these trial-based studies an animal’s actions never affect the situations it will encounter next, a decoupling that is the antithesis of natural foraging, where actions have future consequences, and movement through space changes what the animal can sense and exploit.

Theoretical frameworks provide normative benchmarks for this decision process. The marginal value theorem (Charnov, 1976) predicts when an animal should leave a patch, while the matching law characterizes how choices should be distributed across concurrently available options in proportion to experienced reward rates. Recent computational work has reframed these accounts as inference-and-control problems, in which animals maintain beliefs about latent reward availability and use those beliefs, together with action costs, to guide behavior (Wu et al., 2020; Kilpatrick et al., 2021; Mah et al., 2023). This reframing motivates experimental designs in which the quality of sensory evidence can be parametrically controlled.

Recent studies have addressed this by examining foraging in freely moving primates interacting with real physical environments (Eisenreich et al., 2019; Jacob et al., 2021; Berger et al., 2020; Milton et al., 2020; Voloh et al., 2023; Manea et al., 2024). These studies, together with analogous work in humans (Hutchinson et al., 2008), establish that quality-dependent allocation and systematic, quality-ordered patch transitions are robust features of foraging behavior across species and experimental contexts. However, three features of the foraging problem critical to naturalistic behavior remain unexplored. First, prior tasks have offered animals two simultaneous options, a design in which a single comparative value signal suffices and independent tracking of multiple patches is never required (Louie et al., 2013). Second, the effects of sensory cue reliability have not been systematically quantified: animals either received no cue about reward state or a fully informative one, leaving inference under graded uncertainty unaddressed (Shahidi et al., 2024; Noel et al., 2024). Third, virtually all prior work has used memoryless exponential schedules with constant hazard rates, precluding study of how the temporal structure of reward availability shapes anticipatory behavior and timing (Janssen & Shadlen, 2005).

Here, we developed a dynamic foraging paradigm in which freely moving macaque monkeys navigated a physical arena to harvest rewards from three patches. Rewards became available at random times and then remained available until the monkey pushed a button, at which point reward was delivered and the patch was reset. The availability at each patch evolved independently under stochastic interval schedules and was signaled by a dynamic visual stimulus whose reliability we controlled parametrically. This design allows independent manipulation of temporal hazard structure, cue semantics, and cue reliability, the three dimensions left unexplored by prior work.

We implemented two task variants that differed in reward availability interval statistics and in the quantity encoded by the sensory cue. In one “exponential” variant, reward became available at events generated with a constant hazard rate, giving an exponential distribution of intervals between reward availability; the visual cue conveyed a probabilistic estimate of current reward availability based on patch statistics and elapsed time. In the “gamma” variant, we sampled reward availability events following a time-dependent hazard rate, leading to a gamma distribution of intervals between these events. This reduced the probability that premature responses would be rewarded. We also altered the cue semantics to encode normalized time progression within the currently sampled interval. Within each variant, we manipulated cue reliability across multiple levels.

We found that monkeys allocated pushes according to patch quality, preferentially transitioned toward higher-value options, and progressively refined their allocation over the course of each session. The speed and precision of this allocation relative to patch quality depended on cue reliability to a degree that was substantially amplified under the gamma schedule. These effects demonstrate that temporal reward statistics and sensory uncertainty jointly govern adaptive foraging in continuous environments.

## RESULTS

Macaque monkeys performed a dynamic foraging task designed to approximate natural foraging behavior within a controlled laboratory environment. Animals navigated freely within a hexagonal arena (**Fig. 1A**), moving between three symmetrically positioned foraging patches located on the arena walls (**Fig. 1B**).

**Figure 1.**
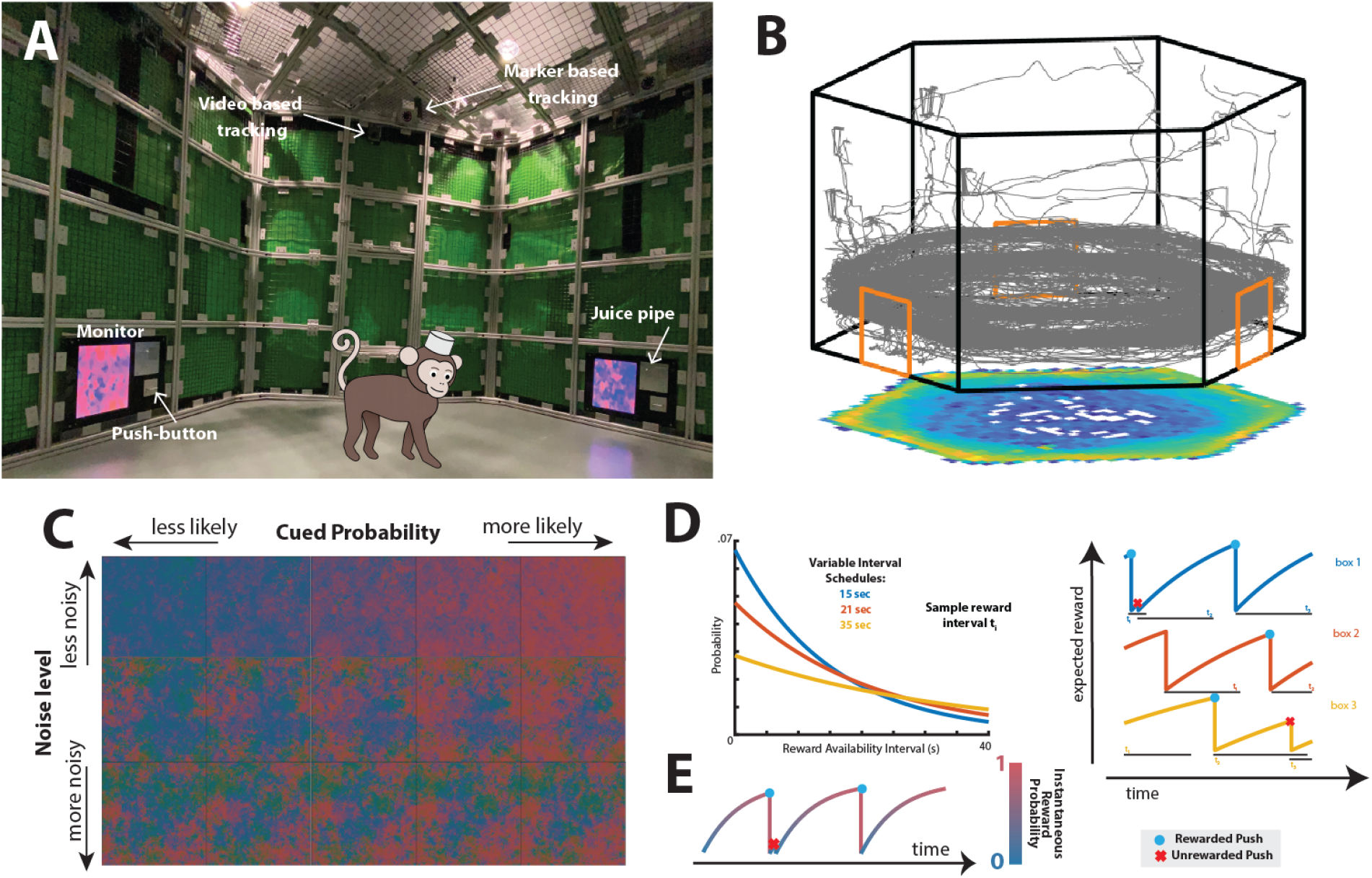
Experimental design to study natural foraging behavior in freely moving monkeys. **A)** The hexagonal arena employed for experiments with freely moving monkeys. **B)** An illustrative trajectory of a monkey performing the foraging task within the freely moving arena. The associated heatmap at the bottom denotes position occupancies. **C)** A stimulus with controllable uncertainty: upon the lever being pushed, the stimulus transitions from blue to progressively more intense shades of red (horizontal). The visual stimulus is scale-invariant (1/f spectrum), thus preventing the subject from gleaning additional information based on the proximity to the stimulus. The reliability of the visual stimulus is modulated by varying the 1/f texture’s variance (vertical). **D)** *Left*: Exponential reward interval distributions from which reward intervals for the three boxes were drawn (blue: fast, orange: medium, yellow: slow). *Right*: Continuous reward availability probability for each box. The horizontal lines at the bottom indicate the currently sampled reward interval. If the button was pressed before the reward interval had elapsed, no reward was delivered and a new interval was sampled (red x). Conversely, if the button was pressed after the reward interval had elapsed, the reward was delivered and the interval was reset (blue dot). **E)** Probability of reward availability for one box during an example segment of the exponential version of the experiment. At each time point, the mean visual stimulus is color-coded by probability. The transition from blue to red thus indicates how soon the reward may become available.

In the exponential variant of the task, rewards at each patch became available according to a variable-interval schedule. Reward availability intervals (RAI) were drawn independently for each box from exponential distributions with fast, medium, and slow rates *µ* (**Fig. 1D, left**). We refer to these rates as determining box “quality”: higher quality means that rewards become available faster. Under this memoryless process, the hazard rate remained constant over time; however, the cumulative probability that reward had already become available increased exponentially as a function of elapsed time since the last press. If a subject pressed the lever after reward became available, a juice reward was delivered, and a new interval was drawn. If the lever was pressed before interval completion, no reward was delivered and the interval was reset (**Fig. 1D, right**).

Reward-related information was conveyed through a dynamic visual stimulus with controllable reliability (**Fig. 1C**). The spatial mean color of this stimulus encoded the probability that reward had already become available by time Δ*t* since the last press, given by the cumulative distribution function of the exponential interval, 1 − *e*^−µ Δ*t*^(**Fig. 1E, Methods**). Thus, at any moment, the cue represented the likelihood that a press would be rewarded, increasing monotonically with elapsed time. To this mean color cue we added a spatiotemporal scale-invariant texture (Field, 1987; Pitkow & Meister, 2012) so subjects would observe the same statistics at all distances from the stimuli. Stimulus reliability was manipulated by varying the variance of this texture. Subjects could therefore use both internal temporal estimates and the externally provided probabilistic cue to guide reward harvesting. Each lever press yielded binary auditory feedback indicating whether the action was rewarded. In this variant, the sensory cue reflected a probabilistic estimate of reward availability derived from patch statistics and elapsed time, rather than information about the specific interval currently in effect.

Across sessions, all three subjects allocated pushes systematically according to patch quality (**Fig. 2A**). Spearman correlations between push fraction and quality rank were positive and highly significant for Monkeys V and M (V: ρ=+0.521, n=597, p<10^−3^; M: ρ=+0.475, n=279, p<10^−3^), and weaker but still significant for Monkey D (ρ=+0.290, n=381, p<10^−3^). At the level of individual blocks, significant positive correlations were obtained in 79%, 80%, and 65% of blocks for Monkeys V, M, and D respectively, confirming consistent matching across sessions.

**Fig 2.**
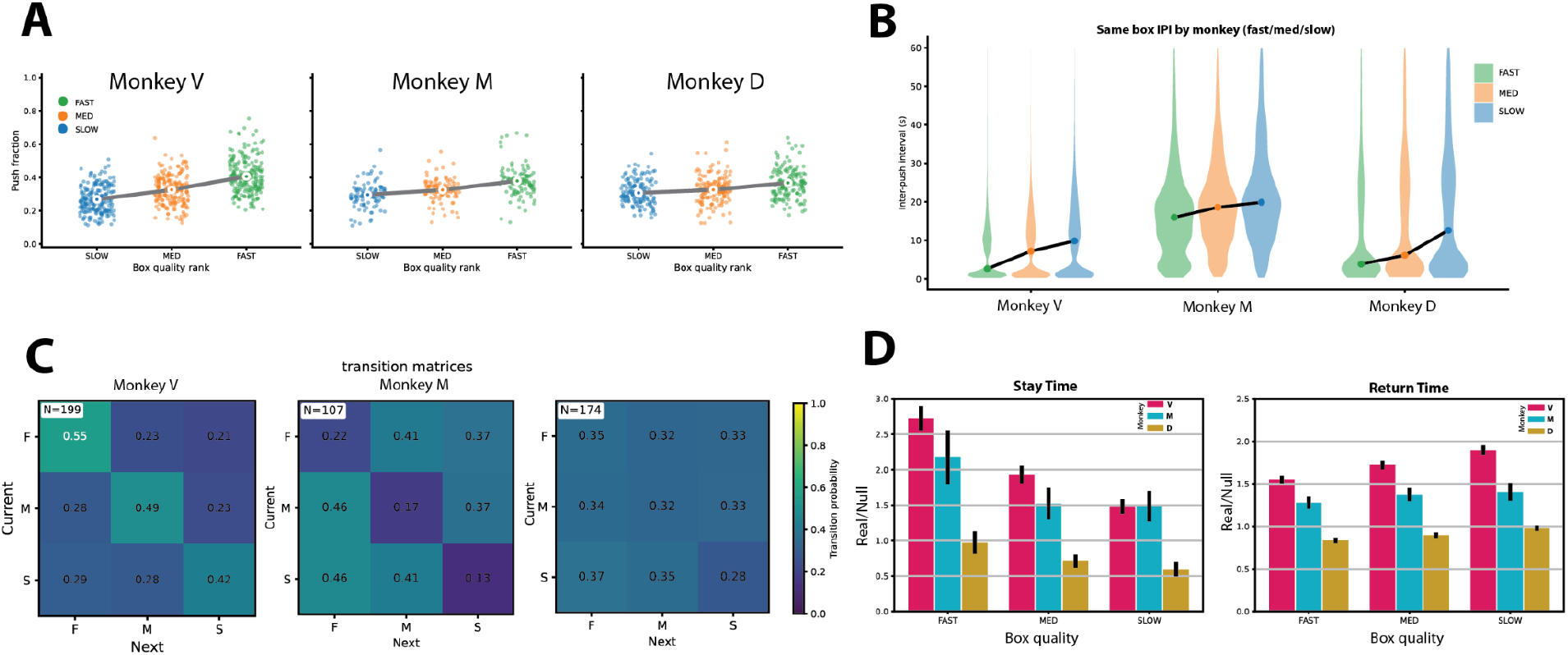
**A)** Push fraction vs. box quality rank for each monkey. Each point is one box observation for one block, color-coded by box type (slow: blue; medium: orange; fast: green). Large dots show mean for each box type; lines connect the means. **B)** Violin plots of inter-push intervals (IPIs) for fast, medium, and slow boxes for each monkey. The central dot within each violin denotes the median inter-push interval. **C)** Transition matrices showing box-switching behavior between fast, medium, and slow boxes. Matrix entries represent the probability of transitioning from the current box (x-axis) to the subsequent box (y-axis), with color indicating transition probability. **D)** Normalized stay times (left) and return times (right) at each box for each monkey. Within each grouped bar plot, bars correspond to box type, and bar color denotes monkey identity. Bar heights indicate the mean across blocks, and error bars represent the SEM across blocks.

The temporal dynamics of within-patch engagement was quantified by inter-push interval (IPI) distributions (**Fig. 2B**). IPIs were bimodal within each box, with short intervals associated with rapid within-patch exploitation and longer intervals associated with a combination of disengagement and between-patch transitions. Median IPIs were shortest at the fast box and longest at the slow box for all three subjects, indicating more persistent local engagement at higher-quality patches.

To characterize switching behavior, we computed transition matrices (**Fig. 2C**), quantifying the probability of remaining at a box or switching to another following a push. Self-transition probabilities were highest at the fast box and decreased monotonically with quality across all subjects. When subjects switched patches, transitions were directed preferentially toward higher-value options. The quality-dependence of transition probabilities was consistent across individuals despite differences in overall persistence levels.

We next quantified stay time and return time (**Fig. 2D**), normalized by a rate-matched null model to control for variation in overall push rate between blocks (see Methods). Stay time scaled systematically with box quality in all subjects, with longest persistence at the fast box and shortest at the slow box. Return time showed the opposite pattern, with faster revisits to higher-value patches. These patterns were robust in Monkeys V and M, and present but weaker in Monkey D.

Together, these results show that subjects systematically organized their push allocation, switching behavior, and temporal patch engagement according to box quality, in a manner consistent with reward-maximizing foraging strategies.

The task included a visual stimulus with controllable uncertainty, allowing us to examine how cue reliability influenced foraging behavior (**Fig. 3**).

**Figure 3.**
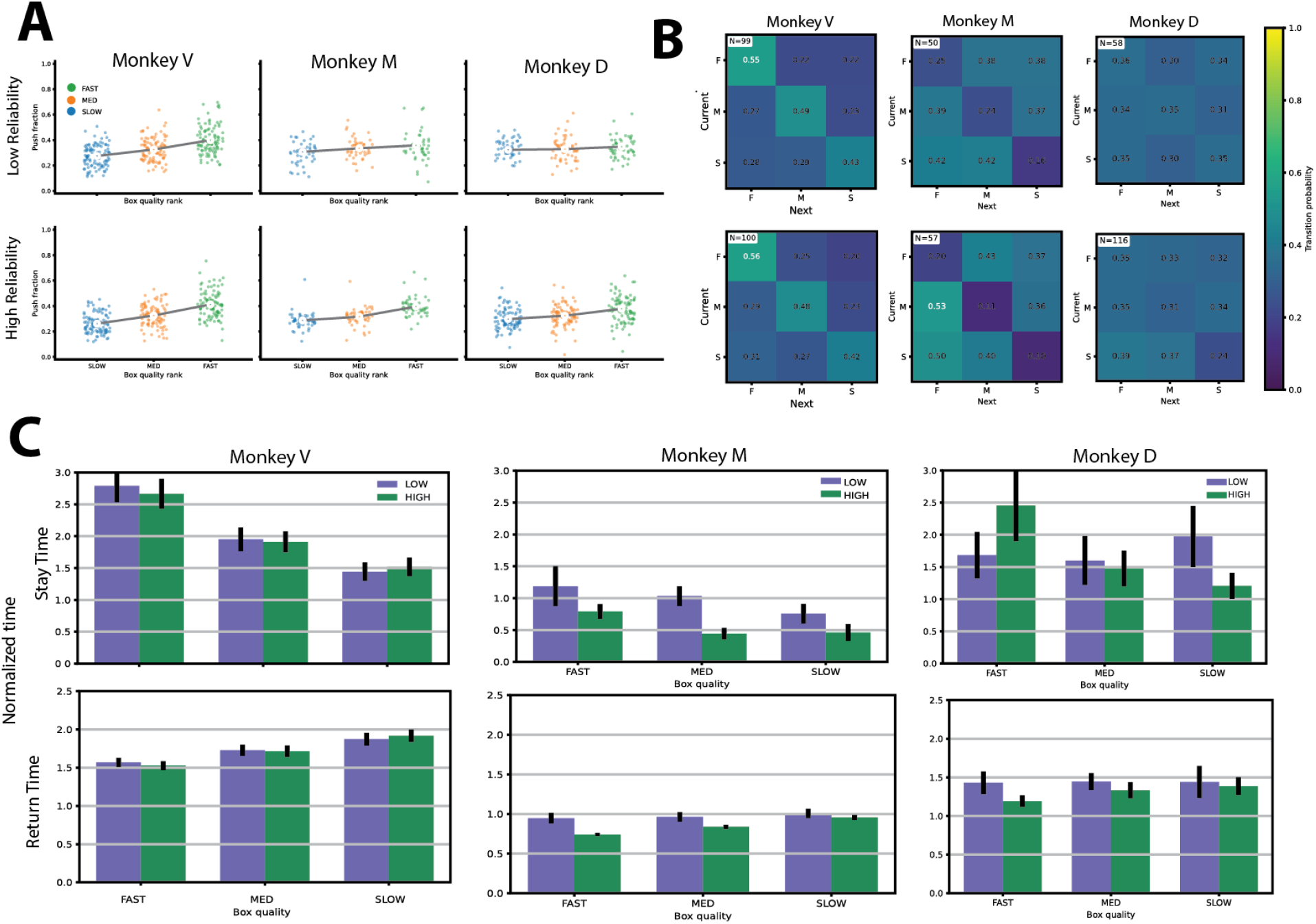
Behavioral performance separated by stimulus reliability. **A**) Push fraction vs. box quality rank for each monkey (columns) split by cue reliability (top row: low reliability; bottom row: high reliability). Each point is one block–box observation, color-coded by box type (slow: blue; medium: orange; fast: green). Large dots show mean per box type; lines connect the means. **B**) Transition matrices illustrating box-switching behavior between fast, medium, and slow boxes. Columns correspond to monkeys; the top row shows low stimulus reliability and the bottom row shows high stimulus reliability. Matrix entries represent the probability of transitioning from the current box (x-axis) to the subsequent box (y-axis), with color indicating transition probability. **C**) Normalized stay times (top row) and return times (bottom row) for each monkey (columns) and each box. Within each grouped bar plot, bars represent box type and color denotes stimulus reliability (green: high reliability; purple: low reliability). Bar heights indicate the mean across blocks, and error bars represent the SEM across blocks.

The degree to which push allocation tracked box quality increased markedly with cue reliability (**Fig. 3A**). Spearman correlations between push fraction and quality rank were consistently higher under high vs. low reliability across all three subjects, suggesting that the monkeys used the visual cues to estimate box quality. For Monkey V, correlations were ρ=+0.461 (n=297, p<10^−3^) under low reliability and ρ=+0.580 (n=300, p<10^−3^) under high reliability. The reliability effect was more pronounced in Monkeys M and D: for Monkey M, ρ rose from +0.202 (n=126, p=0.023) under low reliability to +0.720 (n=153, p<10^−3^) under high reliability; for Monkey D, the low-reliability correlation was not significantly different from zero (ρ=+0.119, n=144, p=0.155), whereas under high reliability it was robust (ρ=+0.390, n=237, p<10^−3^). Thus, while Monkey V maintained meaningful quality-proportional allocation under both conditions, Monkeys M and D required reliable cue information to exhibit consistent preference for higher-value patches.

Transitions between boxes also depended on reliability (**Fig. 3B**). The quality-ordered self-transition pattern was preserved under both conditions in all subjects. For Monkey V, self-transition probabilities were nearly identical across reliability conditions (low: P(F→F)=0.55, P(M→M)=0.49, P(S→S)=0.43; high: 0.56, 0.48, 0.42), indicating robust transition structure regardless of cue reliability. In contrast, Monkey M showed a clear reliability effect in directed switching: the probability of transitioning to the fast box after pushing at another box increased from 0.41 under low reliability to 0.51 under high reliability. Monkey D showed more modest differences in directed switching (0.32 vs. 0.34), despite showing a clear reliability effect in push allocation.

Stay and return time dependence on box quality was largely preserved across reliability conditions for Monkey V (**Fig. 3C**): stay times remained quality-ordered under both reliability conditions, and return times were similarly stable. For Monkey M, the dependence of return time on quality was more apparent under high reliability than low reliability. Monkey D showed weak dependence of stay and return times on box quality under both conditions, with near-flat quality differences particularly under low reliability.

Overall, reducing cue reliability weakened the correspondence between push allocation and box quality and weakened the quality-dependence of patch transitions, but did not abolish quality-dependent behavior. The impact was strongest in Monkeys M and D, where low-reliability conditions approached near-chance allocation, while Monkey V maintained systematic, quality-dependent behavior across both conditions. These individual differences suggest that subjects vary in how much they rely on the sensory cue versus internal estimates of reward availability to guide patch choice.

Sessions lasted approximately 15 minutes, allowing us to examine how the monkeys changed where they allocated their pushes across a block (**Fig. 4A**).

**Figure 4.**
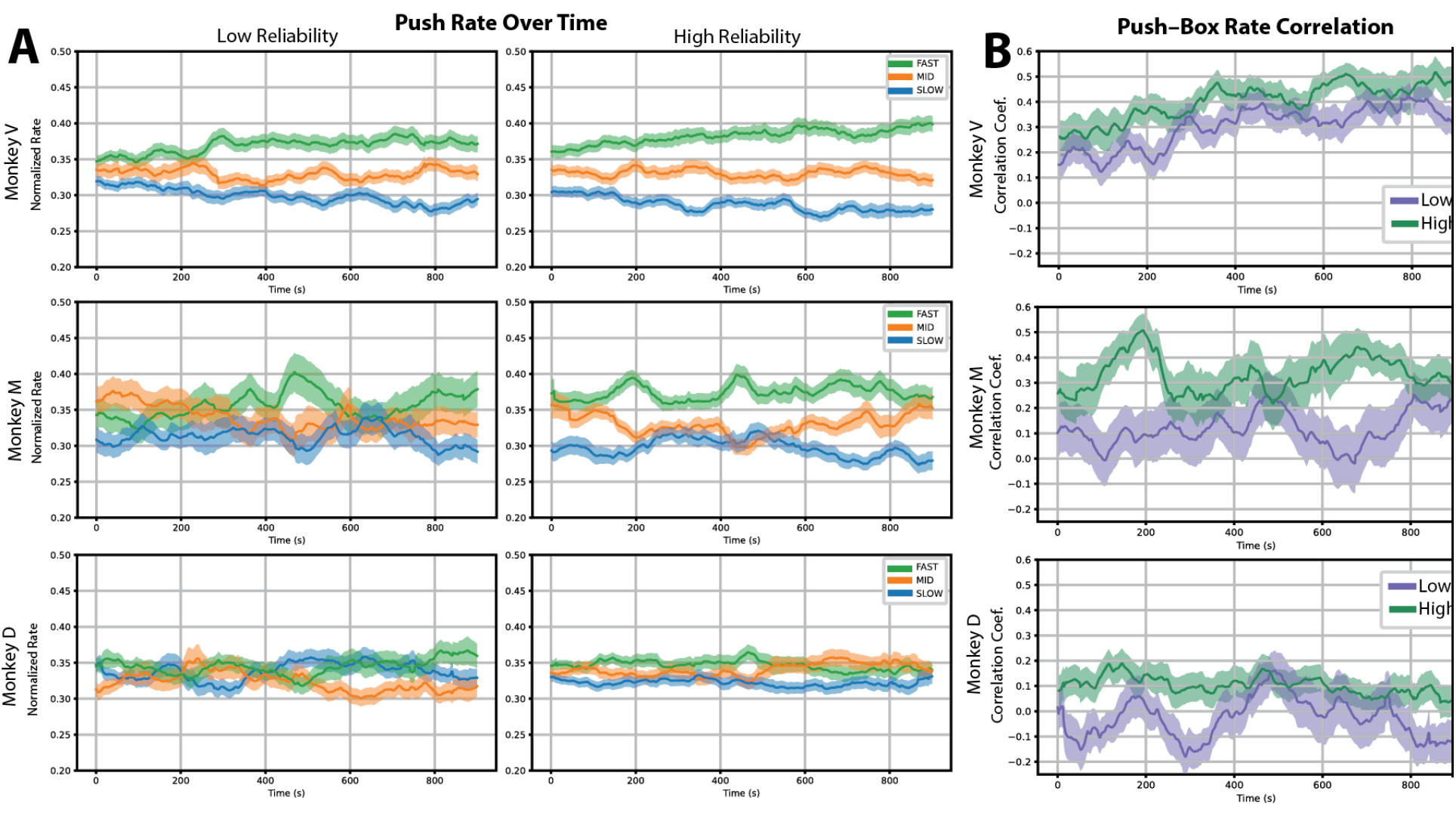
Temporal evolution of push allocation under different stimulus reliability conditions. **A)** Time course of normalized push fraction at each box for low-reliability stimuli (left column) and high-reliability stimuli (right column). Rows correspond to individual monkeys. Push fractions are shown separately for fast, medium, and slow boxes. Solid lines indicate the mean across blocks, and shaded regions represent the SEM across blocks. **B)** Time course of the Spearman correlation coefficient between push fraction and the reward rate parameter across boxes. Correlation values are shown separately for low-reliability (purple) and high-reliability (green) conditions. Rows correspond to individual monkeys. Solid lines indicate the mean across blocks, and shaded regions represent the SEM across blocks.

At the beginning of each block, the monkeys cannot yet know where the fastest box is, but they can learn from their rewarded pushes and from the color cues they see. Indeed, over the course of each session, normalized push fractions for Monkeys V and M became progressively differentiated across box qualities: they pushed more at the fast box and less at the slow box, while the medium box remained intermediate. Monkeys V and M showed progressive differentiation under both reliability conditions, most clearly under low reliability. Monkey D showed minimal consistent learning under either condition.

To quantify how well the monkeys matched their efforts to the rewards, we computed a time-resolved Spearman correlation between push fraction and box quality at each point within the session (**Fig. 4B**). For Monkeys V and M, correlations rose over the course of each session, with higher starting values and steeper increases under high reliability. Monkey M showed the most pronounced reliability effect, with correlations near zero under low reliability but rising substantially under high reliability. Monkey D showed little within-session improvement under either condition.

Together, these results demonstrate that, within a session, Monkeys V and M adapted their effort to align progressively with reward structure; cue reliability modulated both the starting level and the speed of this adaptation. Monkey D showed little evidence of within-session learning under either condition, suggesting a qualitatively different within-block strategy or a longer timescale of adaptation.

Although the exponential-variant task revealed robust quality-dependent allocation and systematic transition dynamics, the influence of cue reliability was comparatively modest. In particular, we observed limited evidence for progressive refinement of allocation within sessions, especially under low reliability. In the exponential variant, the reward process was memoryless and the sensory cue conveyed a probabilistic estimate of reward availability based on patch statistics and elapsed time. This structure may reduce the cost of imprecise timing and offer less benefit from inference under uncertainty.

To examine how temporally predictable reward availability intervals combined with interval-specific progress information influence allocation dynamics, we implemented a new variant of the task for Monkey V (**Fig. 5A–B**). In this variant, we kept the average reward availabilities but changed the reward intervals to follow a gamma distribution (with shape parameter α=10) corresponding to an increasing hazard rate. We also changed the sensory cue to encode time until the next reward becomes available, rather than typical time until reward becomes available. This design introduced more predictable temporal structure, and increased the behavioral cost of responding prematurely.

**Figure 5.**
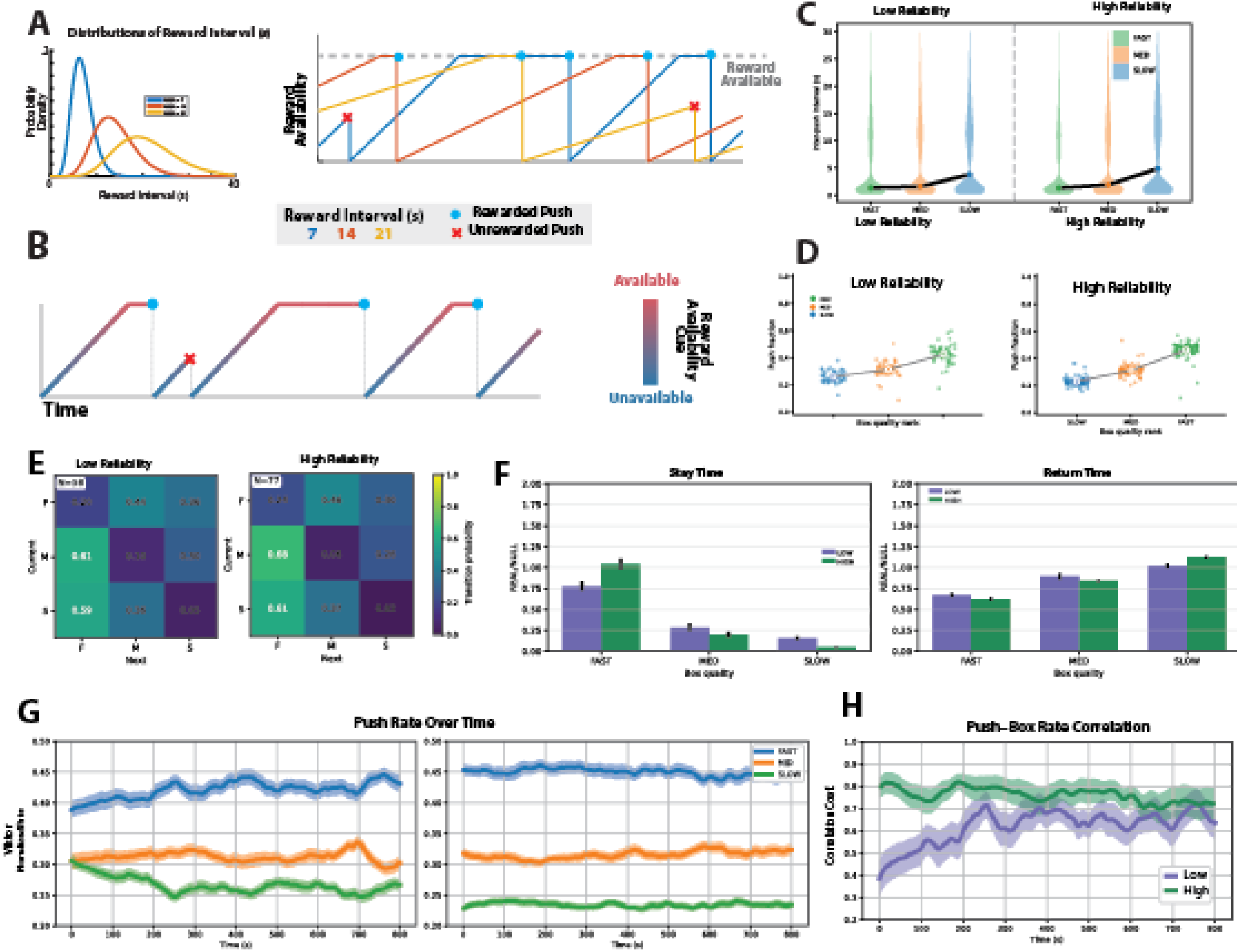
Behavior under gamma-distributed reward schedules with availability-encoding visual cues. **A**) Left: Gamma reward availability interval distributions for the three boxes (blue: fast; orange: medium; yellow: slow). Right: Normalized time progression within the current reward availability interval for each box. Horizontal lines indicate the duration of each sampled reward availability interval. Presses before interval completion yielded no reward and triggered resampling (red ×); presses after interval completion delivered reward and reset the interval (blue dot). **B**) Example reward-availability trace for one box during a representative block, color-coded by the visual stimulus. The transition from blue to red indicates the time until reward becomes available. **C**) Violin plots of inter-push intervals for each box under low-reliability (left) and high-reliability (right) stimulus conditions for monkey V. **D**) Push fraction vs. box quality rank for Viktor, split by cue reliability (left: low reliability; right: high reliability). Each point is one block–box observation, color-coded by box type (slow: blue; medium: orange; fast: green). Large dots show mean per box type; lines connect the means. **E**) Transition matrices for monkey Viktor under low-reliability (left) and high-reliability (right) conditions. Matrix entries represent the probability of transitioning from the current box (x-axis) to the subsequent box (y-axis). **F**) Normalized stay times (left) and return times (right) for monkey Viktor. Dark bars indicate high-reliability cues; light bars indicate low-reliability cues. Bar heights indicate the mean across blocks, and error bars represent the SEM across blocks. **G**) Time course within a representative block of normalized push fraction across boxes under low-reliability (left) and high-reliability (right) conditions. Push fractions are shown for fast, medium, and slow boxes. Solid lines indicate the mean across blocks, and shaded regions represent the SEM across blocks. **H**) Time-resolved Spearman correlation coefficient between push fraction and reward rate parameter under low-reliability (purple) and high-reliability (green) conditions. Solid lines indicate the mean across blocks, and shaded regions represent the SEM across blocks.

Under the gamma schedule, monkeys generated inter-push interval distributions that remained bimodal (**Fig. 5C**), consistent with alternating regimes of within-patch exploitation and between-patch transitions. Median IPIs differed systematically with patch quality (FAST=10.5 s, MED=16.5 s, SLOW=21.2 s), and were substantially longer than in the exponential variant (FAST=1.3 s, MED=1.8 s, SLOW=6.4 s), consistent with the increased cost of consecutive premature pushes under the increasing-hazard schedule.

Push fraction scaled strongly with quality rank under both reliability conditions (**Fig. 5D**): Spearman correlations were ρ=+0.799 (n=168, p<10^−3^) under low reliability and ρ=+0.865 (n=231, p<10^−3^) under high reliability. Both values substantially exceeded the corresponding exponential-variant correlations (+0.461 and +0.580 respectively), indicating that the gamma schedule produced sharper quality-proportional allocation.

Transition dynamics were dramatically reorganized relative to the exponential condition (**Fig. 5E**). Self-transition probabilities were markedly lower overall: P(F→F)=0.23, P(M→M)=0.07, P(S→S)=0.04, compared to 0.55, 0.49, and 0.43 under the exponential schedule. Strikingly, the conditional probability of transitioning to the fast box given a non-fast push reached P(→F|not F)=0.62 — more than double the exponential value of 0.29 — indicating that patch switches were almost predominantly directed toward the highest-value option.

The dependence of stay and return times on box quality was substantially amplified under the gamma schedule (**Fig. 5F**). Under high reliability, stay times dropped steeply with patch quality, with near-zero dwell time at the slow box indicating rapid disengagement after each press. Return times showed the complementary pattern, with active avoidance of the slow box and preferential re-engagement at the fast box. Under low reliability, the same directional pattern held, though with reduced magnitude.

Temporal dynamics within sessions revealed earlier and stronger quality-dependent differentiation of push allocation under the gamma schedule (**Fig. 5G**). Under high reliability, push fractions were already well differentiated between boxes at session onset and remained stable throughout, suggesting near-immediate tracking of reward contingencies based on the timing cue we provided. Under low reliability, where the cue still provided partial information, some differentiation was apparent at session onset on average and continued to develop within the session.

Time-resolved Spearman correlations between push fraction and quality rank corroborated these dynamics (**Fig. 5H**). Under high reliability, correlations were high from session onset and remained stable throughout, indicating that quality-proportional allocation was established near-immediately. Under low reliability, correlations started lower but rose progressively, demonstrating continued within-session refinement even when cue information was degraded. Starting values under both conditions substantially exceeded those observed in the exponential variant.

Overall, introducing temporally predictable reward availability intervals and timing-encoding visual cues strongly amplified both the precision of steady-state allocation and the speed of within-session adaptation. The gamma schedule fundamentally restructured patch transition dynamics — reducing local persistence and concentrating switches toward the highest-value patch — and sharpened the sensitivity of behavior to cue reliability. Together, these results suggest that the temporal statistics of reward availability and cue informativeness jointly shape foraging behavior.

## Discussion

We developed a naturalistic, continuous foraging paradigm that reinstates key ecological properties of real-world decision-making: freely moving navigation, stochastic reward availability, and dynamic sensory cues with controllable reliability. Unlike conventional trial-based paradigms (Sugrue et al., 2004; Corrado et al., 2005; Kolling et al., 2012; Hayden et al., 2011; Blanchard et al., 2015), our task embeds choice within uninterrupted sensorimotor interaction, requiring subjects to continuously integrate internal timing estimates with uncertain external information while allocating behavior across competing reward patches.

Across conditions, monkeys allocated their pushes systematically according to patch quality, consistent with the matching law, exhibited systematic, quality-ordered transition dynamics between patches, and showed quality-dependent scaling of stay and return times. Critically, this quality-proportional allocation did not emerge instantaneously but developed progressively within sessions: push fractions increasingly reflected reward contingencies over time, demonstrating active learning and adaptation within blocks. Beyond average push allocation, monkeys also exhibited temporal structure in how they engaged with patches. Inter-push intervals were bimodal, reflecting alternating regimes of local exploitation and patch switching (Shull et al., 2004), and transitions were directionally biased toward higher-value patches. Together, these findings show that monkeys not only track reward rates but organize behavior temporally and spatially according to box quality, consistent with recent work in freely moving primates (Eisenreich et al., 2019; Jacob et al., 2021; Berger et al., 2020).

Our results demonstrate that monkeys use stimulus information to guide allocation, and that the impact of this information depends jointly on its reliability and the predictability of the reward availability. Under exponential reward availability intervals, monkeys exhibited robust quality-dependent allocation, but the effect of visual reliability was comparatively modest and variable across individuals. The exponential schedule is memoryless: the hazard rate is constant, so elapsed time carries no information about impending reward. Moreover, resetting the availability interval after premature pressing has no impact on the probability distribution of when reward next becomes available, a counterintuitive quirk of the exponential interval schedule; thus early pressing impose only a very limited temporal cost (Janssen & Shadlen, 2005). In this regime, the pressure to precisely track sensory information is reduced, and degraded reliability weakened but did not abolish quality-dependent allocation.

The gamma-distributed reward schedule changed this picture fundamentally. With a time-dependent hazard and an interval-specific cue, reliability effects were substantially amplified. Behavioral differentiation across patches emerged more rapidly within sessions, patch switches were directed preferentially toward the highest-value option, and slow-patch dwell times collapsed to near zero under high reliability. The timing-encoding cue provided a temporally informative signal that mapped directly onto progress within the current reward availability interval. The largest behavioral effect in this study was therefore not the presence of quality-proportional allocation per se, but the interaction between hazard structure and cue reliability: when the environment imposed stronger temporal predictability, sensory reliability had a larger impact on both learning dynamics and the correspondence between push allocation and box quality (Shahidi et al., 2024; Noel et al., 2024). This demonstrates that environmental statistics and perceptual uncertainty jointly govern adaptive behavior

Individual differences were consistent with this framework. All three monkeys exhibited quality-proportional allocation and systematic transition dynamics, but their sensitivity to cue reliability and their degree of within-session refinement varied substantially. Monkeys V and M maintained structured allocation under both reliability conditions, while Monkey D showed near-chance allocation under low reliability and meaningful quality-dependent differentiation of push allocation only under high reliability. Rather than reflecting qualitatively distinct strategies, this variability likely reflects individual differences in how heavily subjects weight sensory evidence relative to internally accumulated reward history (Iigaya et al., 2019), or differences in the timescale over which reward-rate estimates are updated (Sugrue et al., 2004).

Conceptually, foraging in this task can be understood as a dynamic inference problem in which animals estimate latent reward availability under uncertainty (Wu et al., 2020). At each moment, a subject must weigh elapsed time, sensory evidence, knowledge of the schedule structure, and awareness of costs and benefits to decide where to press next. Under the exponential variant, the memoryless schedule and probabilistic cue are consistent with a strategy based on tracking running reward rates across patches. Under the gamma variant, subjects must additionally maintain an estimate of elapsed time within the current interval, a computation that is more sensitive to sensory precision and that compounds the cost of unreliable cue information (Janssen & Shadlen, 2005; Jazayeri & Shadlen, 2010). Future work should formalize these computations using sequential inference models, such as partially observable Markov decision processes (Rao, 2010) or semi-Markov frameworks (Khodadadi et al., 2014; Dayan & Daw, 2008), which can explicitly represent how temporal belief states evolve and how sensory reliability shapes decision boundaries. Such models would allow dissociation of within-session learning dynamics from stable individual traits, and could quantify how subjects weigh internal timing against external sensory evidence across conditions.

More broadly, by manipulating temporal hazard structure and sensory reliability independently in freely moving animals, we demonstrate that adaptive foraging is governed not only by reward rates but by the joint statistics of time, uncertainty, and action cost. This framework establishes a foundation for future neural investigations aimed at uncovering how distributed brain circuits encode hazard structure (Janssen & Shadlen, 2005; Kolling et al., 2012; Hayden et al., 2011), integrate uncertain sensory evidence (Sugrue et al., 2004; Corrado et al., 2005), and continually regulate how they allocate their actions. Previous work using a closed-loop navigation task has shown that sensory, parietal, and prefrontal circuits jointly encode latent task variables during goal-directed behavior (Alefantis et al., 2022; Noel et al., 2022), suggesting that similar distributed representations may support the inference computations required by the present foraging task.

## Material and Methods

### Subjects

#### Monkeys

Three rhesus macaques (male, 6,7 and 8 years old) were used in the experiments. Animal care procedures were conducted in compliance with the National Institute of Health guidelines and were approved by the New York University Animal Care and Use Committee. The animal was chronically implanted with a PEEK headplate (source: Julio Martinez) and an accompanying head cap, CNC-machined from Polyoxymethylene (POM/Delrin) (3D Hubs, USA). The head cap was affixed to the headplate to facilitate head fixation and to protect/support the recording devices

### Experimental set up

Monkeys were tested behaviorally within a hexagonal arena measuring 1.86m in radius and 2.1m in height, constructed from framed panels of 2.54 cm wire mesh (Fig). This design permitted the subjects to move freely in all three dimensions. Three patches were placed on three rotationally symmetric walls of the hexagon. Each patch consisted of a stainless-steel tube connected to a precision valve (Miniature inert liquid valve, Parker Hannifin) for juice delivery, a 15-inch monitor with an integrated speaker (Beetronics, USA, Model: 15VG7M), and a push button (Med Associates Inc.).

The displays on the monitor were rendered using Psychtoolbox 3 in Matlab 2019 (MathWorks). The juice dispensation through the solenoid, managed by an Arduino, was also controlled by MATLAB. Data was collected using the Vicon data acquisition system (Vicon Ltd, UK).

A motion capture system (Vicon Ltd, UK) equipped with 10 infrared Vero cameras was mounted on both the ceiling and wall (Figure S2). This system captured the 3D positions of four reflective markers affixed to the monkey’s head cap at a sampling rate of 100 Hz. The markers were arranged in a single plane on the head cap (Figure): one at the front, one at the back, and the remaining two asymmetrically on the left and right. The 3D coordinates (x,y,z) of each marker were recorded at a rate of 100Hz using Vicon Nexus Software (Vicon Ltd, UK).

The monkey was acclimated to wearing a wireless eye tracking device (ISCAN, Inc., USA). This device was encased in a rigid, 3D-printed shell, which was affixed to the head ring implant. The device featured a miniature infrared camera, an infrared emitter, a hot mirror, and a transmitter. The hot mirror was positioned directly in front of the monkey’s left eye, while the camera was oriented downward at approximately 45 degrees relative to the mirror. For precise 3D head motion tracking, a marker-based motion tracking system was employed. The eye tracking device was powered by a lightweight 3.7 V Li-Po battery and transmitted eye images wirelessly to an external receiver. Both horizontal and vertical eye positions, as well as pupil dilation, were recorded at a frequency of 60 Hz using the Vicon data acquisition system (Vicon Ltd, UK).

Six video cameras (CM3-U3-13Y3C-CS 1/2’’ Chameleon3 Color Camera, FLIR Systems, Inc., USA) were affixed to each wall, capturing footage at 30 Hz from various angles. Additionally, a wide-angle camera was centrally mounted on the ceiling for surveillance purposes.

### Behavioral Task

Subjects performed a continuous three-option dynamic foraging task modeled after a concurrent variable-interval schedule. At any time, each patch generated rewards independently according to a stochastic interval process. Subjects could press a button corresponding to any patch at any moment. If a reward was available at the time of the press, it was delivered; otherwise, no reward was given. In all cases, a new reward interval was sampled immediately following each button press.

Two task variants were implemented. The variants differed in both the distribution governing reward intervals and the variable encoded by the sensory cue. Cue reliability was manipulated independently within each variant by varying the concentration parameter reliability.

#### Task Variants and Informational Structure

The two task variants differ in both temporal reward statistics and cue semantics.

In the exponential variant, reward intervals were memoryless and the sensory cue encoded the cumulative probability of reward availability based on patch statistics and elapsed time since the last press. This mapping provided a probabilistic estimate of availability derived from long-run patch parameters.

In the gamma variant, reward intervals followed a gamma distribution with increasing hazard over time, and the sensory cue encoded normalized progression through the currently sampled reward interval. This mapping provided interval-specific information about progress within the active interval.

Because the encoded quantity differs across variants, comparisons between variants reflect differences in both temporal hazard structure and informational content.

#### Variant 1: Exponential Reward Intervals

Reward intervals for each patch *j*were drawn from an exponential distribution:

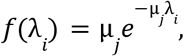

where µ_*j*_is the rate parameter for patch *j*, and the mean interval duration is 1/µ_*j*_. The exponential distribution implies a constant hazard rate:

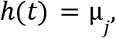

so the instantaneous probability of reward availability is independent of elapsed time since the last press. If a press occurred before the sampled interval elapsed, no reward was delivered and a new interval was sampled (although, counterintuitively, the Markov property implies that a reset is equivalent to maintaining the original interval)

#### Visual Encoding in the Exponential Variant

In this variant, the visual stimulus encoded the cumulative probability that reward was available at time *t* − *t*_*i*_ since the last press:

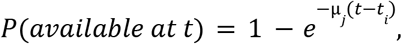

where *t*_*i*_ denotes the time of the most recent press. The cue increased asymptotically over time and reflected a probabilistic estimate of availability derived from elapsed time and patch-specific rate parameters, rather than information about the specific interval currently in effect.

#### Variant 2: Gamma Reward Intervals

In the gamma variant, reward intervals were drawn from a gamma distribution:

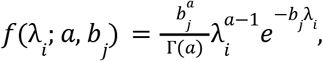

where the shape parameter was fixed at *a* = 10, and *b*_*j*_ was chosen such that the mean interval duration matched that of the corresponding patch. Mean intervals were 7 s, 14 s, and 21 s for the three patches.

For *a* > 1, the gamma distribution produces a time-dependent hazard function that increases over time. This temporal structure increases the cost of premature responding relative to the memoryless exponential case. As in the exponential variant, unrewarded presses triggered a reset and sampling of a new interval.

#### Visual Encoding in the Gamma Variant

In this variant, the visual stimulus encoded normalized progression through the currently sampled interval:

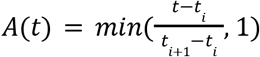

where λ_*i*_ denotes the active sampled interval and *t*_*i*_ is the time of the most recent press.

This variable increased linearly from 0 to 1 as time progressed within the interval and remained at 1 once reward became available. Unlike the exponential variant, the cue therefore provided interval-specific progress information about the active interval rather than a probabilistic estimate derived from long-run patch statistics.

#### Visual Stimulus Construction

In both task variants, reward-related information was conveyed through dynamic naturalistic textures approximating a 1/f power spectrum in spatial and temporal domains.

#### Spatial Spectrum

The spatial amplitude spectrum was defined as:

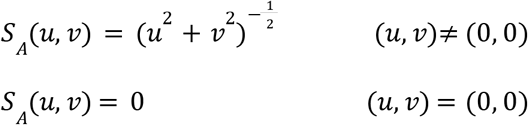

where *u* and *v* are horizontal and vertical spatial frequencies.

#### Temporal Evolution

At each frame *t*, spatial frequency coefficients were updated recursively:

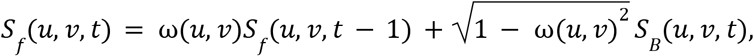

where

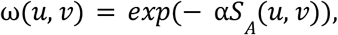

and

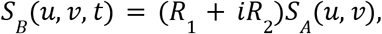

with *R*_1_ and *R*_2_ drawn independently from standard normal distributions.

To ensure a real-valued spatial pattern after the inverse Fourier transform, Hermitian symmetry was enforced:

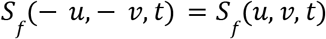

#### Reward-Dependent Modulation

The reward related variable *X*(*t*), representing the version-specific cue signal, was mapped to a phase angle.

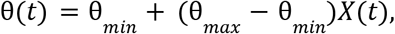

where *X*(*t*) is the cumulative probability of reward availability in the exponential variant and *X*(*t*) = *A*(*t*) in the gamma variant.

The spatial pattern was generated from the spectrum using an inverse Fourier transform (IFFT) as:

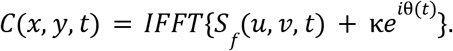

The concentration parameter κ controlled cue reliability. Larger values produced more concentrated phase structure and therefore higher stimulus reliability. Reliability was manipulated within each task variant by varying κ.

#### Color Mapping

The complex spatial pattern was converted to phase angles and mapped to perceptually uniform colors in CIE Luv space via a precomputed lookup table:

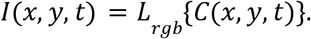

#### Feedback and Reinforcement

For monkeys, rewarded presses yielded a drop of juice and a 440 Hz tone; unrewarded presses produced an 880 Hz tone. For human participants, feedback consisted solely of auditory tones, and successful presses earned points convertible to monetary reward.

### Quantification and data analysis

All data analysis was performed in MATLAB 2021b (The MathWorks, Inc., USA).

#### Extraction of behavioral Variables

Three-dimensional behavioral variables were extracted from the raw marker position data, which underwent pre-processing to account for minor gaps of less than 1 second. All raw data were resampled at a rate of 50 Hz. The position of the head within the horizontal plane was deduced from the x and y components, averaged across all four markers. The amplitude of its time derivative denoted the translational speed. The 3D direction of the monkey’s gaze was determined by merging the eye-in-head position with the 3D head orientation. The spatial view was derived from the intersection between the 3D gaze direction and the surfaces of the hexagonal arena, including walls, floor, and ceiling. In a similar vein, the facing location was gauged from the intersection between the 3D head orientation and these surfaces.

#### Stay and Return Time Analysis

To quantify box engagement dynamics, we computed stay and return times for each box (fast, medium, slow).

**Stay time** was defined as the duration between the first and last push within a contiguous visit to a given box. A visit began with the first push at a box following a switch from another box and ended with the final push before switching away. For visit *i*, stay time was computed as:

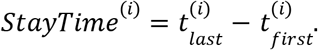

**Return time** was defined as the duration between the last push at a box before leaving and the first push upon returning to that same box:

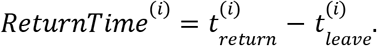

Stay and return times were computed separately for each box identity and pooled across visits within condition.

#### Null Model

To determine whether observed temporal structure exceeded chance expectations given overall push statistics, we constructed a rate-matched null model. Inter-push intervals were drawn from an exponential distribution with rate parameter λ equal to the empirical mean push rate. Each push event was then randomly assigned a box identity (fast, medium, slow), removing structured switching dynamics while preserving overall event rate.

Stay and return times were computed for the null data using the same procedure as for the real data.

#### Normalization

Observed stay and return times were normalized relative to the corresponding values for the null model:

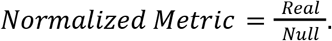

This normalization controls for differences in global push rate and isolates structure attributable to box-specific engagement and switching dynamics.

#### Continuous Push Rate

To characterize the continuous evolution of response vigor over time, we constructed a time-resolved representation of push rate. Let 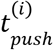 denote the timestamp of the *i*^*th*^ button press, and let

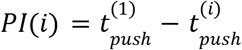

denote the inter-press interval Δ*t*_*i*_ between consecutive presses. We first defined the instantaneous push rate associated with interval *i* as the inverse of the inter-press interval:

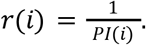

This inversion transforms interval duration into response rate, such that shorter intervals correspond to higher push rates. To obtain a continuous time representation, we constructed a stepwise function over continuous time *t*:

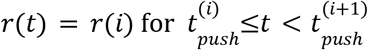

Thus, at any moment between two consecutive button presses, the push rate reflects the rate implied by the most recently completed inter-press interval. This procedure yields a piecewise-constant time series representing ongoing response intensity.

To facilitate comparisons across sessions and subjects, the continuous push rate was normalized at each time frame. Specifically, rates were rescaled relative to the distribution of rates within each session, producing a dimensionless normalized push rate time series.

Finally, to emphasize slower fluctuations in behavioral strategy and reduce high-frequency variability, the normalized push rate was smoothed using a 60-second moving average window.

## Acknowledgements

We thank Zhe Li for feedback on task design and conceptual framing of the analysis. This work was supported by the National Institutes of Health (NIH RF1 NS127122).

## References

Alefantis, P., Lakshminarasimhan, K., Avila, E., Noel, J.-P., Pitkow, X., & Angelaki, D.E. (2022). Sensory evidence accumulation using optic flow in a naturalistic navigation task. The Journal of Neuroscience, 42(27), 5451–5462.

Baum, W.M. (1974). On two types of deviation from the matching law: bias and undermatching. Journal of the Experimental Analysis of Behavior, 22(1), 231–242.

Berger, M., Shahryar Agha, N., & Gail, A. (2020). Wireless recording from unrestrained monkeys reveals motor goal encoding beyond immediate reach in frontoparietal cortex. eLife, 9, e51322.

Blanchard, T.C., & Hayden, B.Y. (2015). Monkeys are more patient in a foraging task than in a standard intertemporal choice task. PLoS ONE, 10(2), e0117057.

Charnov, E.L. (1976). Optimal foraging, the marginal value theorem. Theoretical Population Biology, 9(2), 129–136.

Corrado, G.S., Sugrue, L.P., Seung, H.S., & Newsome, W.T. (2005). Linear-nonlinear-Poisson models of primate choice dynamics. Journal of the Experimental Analysis of Behavior, 84(3), 581–617.

Davison, M., & McCarthy, D. (1988). The Matching Law: A Research Review. Hillsdale, NJ: Lawrence Erlbaum Associates.

Dayan, P., & Daw, N.D. (2008). Decision theory, reinforcement learning, and the brain. Cognitive, Affective, & Behavioral Neuroscience, 8(4), 429–453.

Eisenreich, B.R., Hayden, B.Y., & Zimmermann, J. (2019). Macaques are risk-averse in a freely moving foraging task. Scientific Reports, 9, 15091.

Field, D.J. (1987). Relations between the statistics of natural images and the response properties of cortical cells. Journal of the Optical Society of America A, 4(12), 2379–2394.

Pitkow, X., & Meister, M. (2012). Decorrelation and efficient coding by retinal ganglion cells. Nature Neuroscience, 15(4), 628–635.

Hayden, B.Y., Pearson, J.M., & Platt, M.L. (2011). Neuronal basis of sequential foraging decisions in a patchy environment. Nature Neuroscience, 14(7), 933–938.

Herrnstein, R.J. (1961). Relative and absolute strength of response as a function of frequency of reinforcement. Journal of the Experimental Analysis of Behavior, 4(3), 267–272.

Houston, A.I., Trimmer, P.C., & McNamara, J.M. (2021). Matching behaviours and rewards. Trends in Cognitive Sciences, 25(5), 403–414.

Hutchinson, J.M.C., Wilke, A., & Todd, P.M. (2008). Patch leaving in humans: can a generalist adapt its rules to dispersal of items across patches? Animal Behaviour, 75(4), 1331–1349.

Iigaya, K., Ahmadian, Y., Sugrue, L.P., Corrado, G.S., Loewenstein, Y., Newsome, W.T., & Fusi, S. (2019). Deviation from the matching law reflects an optimal strategy involving learning over multiple timescales. Nature Communications, 10, 1466.

Jacob, G., Katti, H., Cherian, T., Das, J., Zhivago, K.A., & Arun, S.P. (2021). A naturalistic environment to study visual cognition in unrestrained monkeys. eLife, 10, e63816.

Janssen, P., & Shadlen, M.N. (2005). A representation of the hazard rate of elapsed time in macaque area LIP. Nature Neuroscience, 8(2), 234–241.

Jazayeri, M., & Shadlen, M.N. (2010). Temporal context calibrates interval timing. Nature Neuroscience, 13, 1020–1026.

Khodadadi, A., Fakhari, P., & Busemeyer, J.R. (2014). Learning to maximize reward rate: a model based on semi-Markov decision processes. Frontiers in Neuroscience, 8, 101.

Kilpatrick, Z.P., Davidson, J.D., & El Hady, A. (2021). Uncertainty drives deviations in normative foraging decision strategies. Journal of the Royal Society Interface, 18(180), 20210337.

Kolling, N., Behrens, T.E.J., Mars, R.B., & Rushworth, M.F.S. (2012). Neural mechanisms of foraging. Science, 336(6077), 95–98.

Louie, K., Khaw, M.W., & Glimcher, P.W. (2013). Normalization is a general neural mechanism for context-dependent decision making. Proceedings of the National Academy of Sciences, 110(15), 6139–6144.

Mah, A., Schiereck, S. S., Bossio, V., & Constantinople, C. M. (2023). Distinct value computations support rapid sequential decisions. Nature Communications, 14, 7573.

Manea, A.M.G., Maisson, D.J.-N., Voloh, B., Zilverstand, A., Hayden, B.Y., & Zimmermann, J. (2024). Neural timescales reflect behavioral demands in freely moving rhesus macaques. Nature Communications, 15, 2151.

Milton, R., Shahidi, N., & Dragoi, V. (2020). Dynamic states of population activity in prefrontal cortical networks of freely-moving macaque. Nature Communications, 11, 1948.

Noel, J.-P., Balzani, E., Avila, E., Lakshminarasimhan, K.J., Bruni, S., Alefantis, P., Savin, C., & Angelaki, D.E. (2022). Coding of latent variables in sensory, parietal, and frontal cortices during closed-loop virtual navigation. eLife, 11, e80280.

Noel, J.P., Balzani, E., Savin, C., & Angelaki, D.E. (2024). Context-invariant beliefs are supported by dynamic reconfiguration of single unit functional connectivity in prefrontal cortex of male macaques. Nature Communications, 15, 5738.

Rao, R.P.N. (2010). Decision making under uncertainty: a neural model based on partially observable Markov decision processes. Frontiers in Computational Neuroscience, 4, 146.

Shahidi, N., Franch, M., Parajuli, A., Schrater, P., Wright, A., Pitkow, X., & Dragoi, V. (2024). Population coding of strategic variables during foraging in freely moving macaques. Nature Neuroscience, 27, 772–781.

Shull, R.L., Grimes, J.A., & Bennett, J.A. (2004). Bouts of responding: the relation between bout rate and the rate of variable-interval reinforcement. Journal of the Experimental Analysis of Behavior, 81(1), 65–83.

Sugrue, L.P., Corrado, G.S., & Newsome, W.T. (2004). Matching behavior and the representation of value in the parietal cortex. Science, 304(5678), 1782–1787.

Voloh, B., Maisson, D.J.-N., Lopez Cervera, R., Conover, I., Zambre, M., Hayden, B.Y., & Zimmermann, J. (2023). Hierarchical action encoding in prefrontal cortex of freely moving macaques. Cell Reports, 42, 113091.

Wu, Z., Kwon, M., Daptardar, S., Schrater, P., & Pitkow, X. (2020). Rational thoughts in neural codes. Proceedings of the National Academy of Sciences, 117(47), 29311–29320.

